# Identification of *YAB11-NGAL1* controlling leaf serrations through enhanced genome-wide association studies of *Populus*

**DOI:** 10.1101/2022.04.07.487505

**Authors:** Peng Liu, Chenhao Bu, Panfei Chen, Deqiang Zhang, Yuepeng Song

## Abstract

Leaf margins are complex plant morphological features and contribute to the diversity of leaf shapes which effect on plant structure, yield and adaptation. Although several regulators of leaf margins have been identified, the genetic basis of natural variation therein has not been fully elucidated. We first profiled two distinct types (serration and smooth) of leaf morphology using the persistent homology mathematical framework (PHMF) in poplar. Combined genome-wide association studies (GWAS) and expression quantitative trait nucleotide (eQTN) mapping to create a module of leaf morphology controlling using data from *Populus tomentosa* and *P. simonii* association population, respectively. Natural variation of leaf margins is associated with transcript abundances of *YABBY11* (*YAB11*) in poplar. In *P. tomentosa*, *PtoYAB11* carries premature stop codon (*PtoYAB11^PSC^*) resulting in lost its positive regulation in *PtoNGAL-1*, *PtoRBCL*, *PtoATPA*, *PtoATPE*, and *PtoPSBB*. Overexpression of *PtoYAB11^PSC^* serrated leaf margin, enlarged leaves, promoted photosynthesis and increased biomass. Overexpression of *PsiYAB11* in *P. tomentosa* could rescue leaf margin serration and increase stomatal density and light damage repair ability. In poplar*, YAB11*-*NGAL1*is sensitive to environmental conditions and play positive regulator of leaf margin serration. It might be important regulator which bridge environment signaling to leaf morphological plasticity.

## Introduction

Leaf shape has a strong influence on plant structure, fruit yield and quality (Murchie et al., 2009; Rowland et al., 2020). Wider and thinner leaves can promote plant photosynthesis and gas exchange (Tsukaya, 2006). In terms of ecological functions, leaf shape also prevents leaf damage by insects (Higuchi and Kawakita, 2019). Therefore, optimizing leaf shape is a major challenge in breeding ideal plants. Leaves of eudicots exhibit a staggering diversity of forms, including smooth, serrated (elaborated to form shallow protrusions), and lobed (deeper protrusions) forms (Nikolov et al., 2019). Diverse shapes of eudicot leaves result from small variations of three interwoven processes including the patterning of serrations, lobes and/or leaflets on the leaf margin; the patterning of the vascular system; and the growth of the leaf blade spanning the main veins (Runions et al., 2017). The establishment of leaf margins depends on delimitation of the adaxial and abaxial sides of the leaf relative to the central axis. In the Arabidopsis leaf epidermal cell, expression of *PIN-FORMED1* (*PIN1*) is required for the formation of auxin maxima. Together, auxin maxima and the *CUP-SHAPED COTYLEDON2* (*CUC2*) form an alternating pattern along the proximal margin of leaves that is necessary for normal serration patterns (Nikovics et al., 2006). Auxin is also active in regulation of miR164a, a negative regulator of *CUC2* in the establishment of alternating leaf margin patterns (Bilsborough et al., 2011). *NGATHA-LIKE1–3* (*NGAL1–3*) also negatively regulate leaf margin serration formation through modulating *CUC2* transcript abundance (Shao et al., 2020). In addition to the auxin response regulator, *REDUCED COMPLEXITY* (*RCO*) also controls the final shape of protrusions by promoting growth repression near the leaf base (Vlad et al., 2014). This implies that a multi-layered regulatory network is involved in leaf margin development.

Quantitative trait locus (QTL) mapping, linkage mapping, and genome-wide association studies (GWAS) have proven to be powerful tools for understanding complex traits. In *P. trichocarpa*, single nucleotide polymorphisms (SNPs) within *GALACTURONOSYL-TRANSFERASE 9* (*GAUT9*) are significantly associated with leaf length (LL), leaf width (LW), leaf area (LA), and the length-to-width ratio (Chhetri et al., 2019). *PtrYABBY3*, a plant-specific TF linking adaxial–abaxial polarity with leaf growth activity, is significantly associated with LA (Chhetri et al., 2019). LA, LL, LW and the length-to-width ratio can be determined using manual or digital measurements, whereas more complex phenotypes, like leaf margins, need to be defined algorithmically (Biot et al., 2016). To future determine the main regulators of leaf margin nature variation, high throughput and accurate leaf morphological feature analysis is needed.

LAMINA software provides an efficient and accurate means of analyzing LA in large datasets (Bylesjö et al., 2008). MowJoe converts leaf data into graphs through image skeletonization, to yield information on leaflets and leaf shape (Failmezger et al., 2018). Elliptical Fourier descriptors (EFDs) have been used for global analysis of grape leaf outlines and lobe positions (Chitwood et al., 2015). Persistent homology mathematical framework (PHMF), a newly developed topology data analysis method, takes the pixels of the leaf contour as a 2D point cloud, and then applies a neighborhood density estimator to each pixel. Integrating complex plant morphological features into a single descriptor can capture morphological variation more comprehensively than measuring univariate features (Li et al., 2018). These studies indicate that integrating high throughput, accurate leaf morphological feature analysis and GWAS offers a powerful systems genetics tool for identifying the major regulators of leaf margin.

Although several leaf morphology regulators have been identified, environmental influences on leaf shape should not be ignored. Heterophylly is a striking feature of plants that grow at the border of terrestrial and aquatic environments and often withstand partial submergence conditions (Nakayama et al., 2017). Gibberellic acid (GA), abscisic acid (ABA), ethylene, and auxin have also been found to play important roles in the interplay between leaf development and the environment (Nakayama et al., 2017; He et al., 2020; Zhang et al., 2020). In *P. trichocarpa*, *PtrYAB3* was not only associated with leaf morphology, but also had significant effects on stomatal density and carbon isotopes (Chhetri et al., 2019). Understanding the genetic basis of leaf morphology development might also shed light on its adaptive significance.

*Populus* consist of five subgenera including aigeiros, leuce, leucoides tacamahaca and turanga contain abundant leaf morphological diversity within and among different poplar species which is an ideal model plant for analysis of genetic basis of natural variation of leaf type (Song et al., 2020). In this study, PHMF was used to accurately characterize the two distinct type leaf morphological features of a natural population of *P. tomentosa* and *P. simonii*. Integrated GWAS and eQTNs results showed that *YAB11*-*NGAL1* is a major regulator of the margins of *Populus* leaves. Overexpression of *PtoYAB11^PSC^* led to serrated leaf margins, enhanced photosynthesis, and increased biomass in *P. tomentosa*. In contrast, overexpression of *PsiYAB11* smoothed leaf margin and increased stomatal density, iWUE and light damage repair ability. Both of *PtoYAB11^PSC^* and *PsiYAB11* expression were sensitive to environmental signaling. This study introduces a novel molecular mechanism involved in leaf morphological development that responds to environmental conditions.

## Materials and methods

### Plant materials for DNA and RNA extraction

A total of 940 unrelated 11-years-old *Populus* accessions including 435 *P. tomentosa* and 505 *P. simonii* unrelated individuals provided the material for GWAS and eQTN mapping. The 940 accessions (three biological repetitions per genotype) were grown in a clonal arboretum with three clonal replicates planted according to a randomized complete block design (3 × 4 m spacing). One-year-old *P. tomentosa* clone ‘1316’ with the same height (∼50 cm) were subjected to abiotic stress and phytohormone treatments, and tissue-specific gene expression analysis (Supplemental Methods S1). Leaf plastochron index (LPI) were used to distinguish phase of leaf development (Taylor et al., 2003). For *P. tomentosa* and *P. simonii* population expression variation analysis, mature leaves (LPI 7) of all samples were collected and frozen in liquid nitrogen. Total RNA all of frozen samples were extracted using the Qiagen RNAeasy kit following the manufacturer’s protocols.

### Persistent homology mathematical framework for leaf morphology analysis

Five mature leaves (LPI7) which sampled from each of three replicate clonal copies of each genotype were individually scanned using the CanoScan LIDE300 instrument (Canon, Tokyo, Japan). Binary images of the leaves were made using a Python script (Available on request). Elliptical fourier descriptors (EFDs) were used for approximation of leaflet contours (Iwata and Ukai, 2002). EFDs decompose the contour into a weighted sum of wave functions with different frequencies (Supplemental Methods S2). The euler characteristic and EFD approximation were used to quantify leaflet serrations. PHMF analyses were performed using MATLAB software (MathWorks, Inc., Natick, MA, USA), and the R (R Development Core Team, Vienna, Austria) packages factoextra and FactoMine R package were used for principal component analysis (PCA) mapping (Kassambara and Mundt, 2020). Image J software (NIH, Bethesda, MD, USA) was used to measure traditional leaf shape parameters including leaf length (LL), leaf width (LW), leaf area (LA), leaf perimeter (LP), leaf aspect ratio (AR), leaf area perimeter ratio (AP), and leaf perimeter length ratio (PL).

### GWAS of PHMF leaf morphology-related traits

SNP heritability (*H^2^*) for each phenotype was estimated using the multiBLUP tool (Speed et al., 2014). All of 945 *Populus* accessions (N_Pto_ = 435; N_Psi_=505)of the association population were sequenced with an average of 15× (raw data) coverage using the Illumina GA2 sequencing platform. The Clean reads were mapped to the *Populus* reference genome v3.0 and the SNP calling pipeline were executed as described previously (Xiao et al., 2019). In total, 2,757,176 and 1,305,980 SNPs (MAF >5% and missing rate < 20%) were derived, respectively. The mixed linear model (MLM) was implemented in TASSEL 5.2 software (Bradbury et al., 2007) and used to analyze associations between whole-genome SNPs and leaf morphology-related traits. To control for possible confounding effects of population structure (Q) and pairwise kinship (K), both were included in the mixed linear model. The optimal substructure result (K_Pto_=3; K_psi_=4) was assessed using a statistical model described by Du et al., (2012) and Song et al (2020). *P*-values were calculated, with significance defined as *P*_Pto_ ≤ 1.81E – 08 (*P*<0.05/n, N_Pto_=2,757,176), *P*_Psi_ ≤ 7.65E – 08(*P*<0.05/n, N_Psi_= 1,305,980) based on Bonferroni correction. Sanger sequencing were used to validate candidate SNPs (Supplemental Methods S3).

### RNA-sequencing and eQTNs mapping

All of 945 poplar accessions leaves (LPI 7) cDNA library construction and sequencing were performed by Beijing Biomarker Technology (Beijing, China) (Supplemental Methods S4). Raw sequencing reads were mapped to *P. trichocarpa* reference genome v3.0 genome using HISAT2 (Kim et al., 2015). The expression levels of fragments per kilobase of transcript per million fragments (FPKM) were calculated using Cuffquant and Cuffnorm software (Trapnell et al., 2012). Transcripts expressed in more than 80% of two population were selected for eQTNs analysis (Supplemental Methods S5). The linear regression mode of the Matrix eQTNs Package was used to identify associations between SNPs derived from the *PtoYAB11^PSC^* and *PsiYAB11* genes (± 2,000 bp flanking region) and the expression levels of genome-wide genes (Shabalin, 2012). Bonferroni correction was applied for multiple comparisons (*P* < 1.0E-05) to control for genome-wide false-positives. All eQTNs located within a 10^4^ bp area around the transcription start site (TSS) of targets were regarded as *cis*-eQTNs, while all other eQTNs were regarded as *trans*-eQTNs (Quan et al., 2019).

### Poplar *YABBY* (*YAB*) gene family identification and phylogenetic analysis

All *YAB* gene family member sequences were cloned from *P. tomentosa* ‘1316’ and *P. simonii* ‘QL9’. *YAB11* gene sequences were also cloned from *P. simonii*, *P. nigra*, *P. szechuanica*, and *P. euphratica*, respectively. The amino acid sequences of the Arabidopsis and rice (*Oryza sativa*) *YABBY* gene families were acquired from Phytozome (https://phytozome.jgi.doe.gov/) (Supplemental Methods S6). Clustal X 2.0 (Larkin et al. 2007) and MUSCLE (Edgar, 2005) software were used to align the YABBY protein sequences, to identify YABBY and zinc-finger domains. For phylogenetic analysis, Clustal W (Kumar et al., 2016) was used to align the full-length YABBY sequences from rice. For Arabidopsis, rice and poplar, a maximum likelihood (ML) tree was constructed using the bootstrap method with 1,000 replicates. The minimum evolution method was used to validate the ML tree.

### Subcellular localization

The full-length coding sequences of *PsiYAB11* and *PtoYAB11^PSC^* was cloned into the pBI121 vector to generate green fluorescent protein (GFP) fusion constructs driven by 35S:promoter. Then, 35S: *PsiYAB11*-GFP and 35S: *PtoYAB11^PSC^*-GFP were transiently expressed in *Nicotiana benthamiana* leaves. The fluorescence signal was observed in *N. benthamiana* leaves (Three biological repetition) with confocal laser scanning microscopy (C2-ER; Nikon, Tokyo, Japan) two days after transformation (Supplemental Methods S7).

### DNA Affinity Purification sequencing (DAP-seq) and data processing

Five μg *P. tomentosa* genomic DNA (gRNA) was fragmented to an average of 200 bp and used for DNA-seq library construction. PsiYAB11 protein was expressed using the TNT SP6 Coupled Wheat Germ Extract System (Promega). The protein-bound beads were incubated with 50 ng of adaptor-ligated gDNA fragments on a rotator for 1 h at room temperature. After denature the protein and release the bound DNA fragments, eluted DNA fragments were sequenced on an Illumina NavoSeq. Reads were mapped to *P. trichocarpa* reference genome v3.0 using HISAT2 (Kim et al., 2015). DAP-seq peaks, which stand for the transcription factor binding sites, were called by MACS2 coupled with the IDR pipeline (Zhang et al., 2008). Motif discovery was performed using MEME-Chip suite 5.0.5 (Machanick et al., 2011) (Supplemental Methods S8).

### Stomatal densities and size measurement

Stomatal morphological characteristics were measured by scanning electron microscopy (Hitachi S-3400 N II, Tokyo, Japan). Two small pieces were cut from the leaves from each genotype between the midrib and the leaf margin. One piece was used for adaxial observation and another piece was used for abaxial observation. Observations were made at ×320–4000 magnification. Five randomly selected fields of the leaf surface were recorded. Stomata counting and density calculation were performed as described by Pearce et al. (2005) (Supplemental Methods S9).

### Transient transcription dual-luciferase (LUC) assay

The *PtoNGAL1*, *PtoCUC2*, *PtoRBCL*, *PtoATPA*, *PtoATPE* and *PtoPSBB* promoter was amplified from *P. tomentosa* ‘1316’ respectively and cloned into the modified pGreenII 0900 vector containing the firefly luciferase (fLUC) gene and the Renilla luciferase gene (rLUC) as reporters, while the *PsiYAB11*, *PtoYAB11^PSC^* and *PtoNGAL1* cDNAs, were cloned into the pBI121 expression vector as an effector. Plasmids were transferred into *A. tumefaciens* GV3101 by electroporation and coin filtrated into *N. benthamiana* leaves. Luciferase activities were measured using the Dual Luciferase Reporter Assay System (Promega, Madison, WI, USA) according to the manufacturer’s instructions. Relative reporter gene expression levels were expressed as the ratio of fLUC to rLUC. Six independent transformations of each sample were performed (Supplemental Methods S10).

### Yeast one-hybrid (Y1H) assays

The *PsiYAB11* and *PtoYAB11^PSC^* genes were cloned into the pGADT7-rec vector (Clontech, Mountain View, CA, USA) with the GAL4 activation domain. The promoters of *PtoNGAL1*, *PtoNGAL1*, *PtoCUC2*, *PtoRBCL*, *PtoATPA*, *PtoATPE* and *PtoPSBB* were inserted into a pAbAi-BR yeast-integrating vector (Clontech), which was used as a bait-reporter construct. pGADT7-*PsiYAB11*, pGADT7-*PtoYAB11^PSC^*, and pAbAi-^Pro^ (*PtoNGAL1*, *PtoNGAL1*, *PtoCUC2*, *PtoRBCL*, *PtoATPA*, *PtoATPE* and *PtoPSBB*) were co-transformed into the Y1HGold yeast strain using the Matchmaker One-Hybrid Library Construction and Screening Kit (Clontech), respectively. Co-transformed yeast cells were selected on synthetic dextrose minimal medium (SD) lacking uracil (Ura) and leucine (Leu) (SD/-Ura-Leu), with or without 0.6 mg/L Aureobasidin A (AbA, Sigma-Aldrich, St. Loui, MO, USA), and were then incubated for 5 days at 30°C. pGAD-p53+p53-pAbAi and pGADT7-AD+ pAbAi-^Pro^ were carried out in the same manner as for the positive and negative controls, respectively. The primers used for these analyses are listed in Supplemental Table S1 (Supplemental Methods S11).

### Overexpressed PsiYAB11 and PtoYAB11^PSC^ in P. tomentosa

The coding sequences of *PsiYAB11* and *PtoYAB11^PSC^* were cloned into the pBI121 vector driven by a 35S promoter and introduced into Agrobacterium GV3101 (Supplemental Figure S1). All primers for plasmid construction and quantitative polymerase chain reaction (qPCR) are listed in Supplemental Table S1. Agrobacterium, with the overexpressed vector, was used for infection of *P. tomentosa* ‘1316’ leaf discs. Overexpressed transgenic *P. tomentosa* leaf discs were selected on hygromycin-containing (50 mM) medium. Six transgenic *P. tomentosa* shoot were cut into 0.2–0.3 cm pieces and transferred into a proliferation medium (Murashige and Skoog [MS] + 2 mg/L 2,4-D + 1 mg/L 6-BA + 30 g/L sucrose + 4.5 g agar). Data were obtained from at least three transgenic lines and WT that showed stable phenotype. Height, ground dimeter, and leaf morphology traits of all samples were measured after transfer into plot 120 days (Supplemental Methods S12).

### Chlorophyll fluorescence-based probes of photosynthetic parameters

The sixth fully expanded leaf of transgenic *P. tomentosa* ‘1316’ were used for photosynthetic rate measurements and chlorophyll fluorescence analysis. Three independent biological clones were analyzed for each genotype. Photosynthetic rates were measured using a portable photosynthesis system (LI-6400XT; Li-Cor Biosciences, Lincoln, NE, USA) on 15 August 2020 from 9:00 am to 11:00 am. The net photosynthetic rate (Pn), transpiration rate (Tr), intercellular CO_2_ concentration (Ci), and stomatal conductance (Gs) were measured simultaneously. Intrinsic water use efficiency (iWUE) was calculated from the ratio of Pn to Tr. Chlorophyll fluorescence parameters were estimated by the MultispeQ instrument (Photosynq, East Lansing, MI, USA) using pulse amplitude modulation (PAM) fluorometry (Baker et al., 2007). Variable fluorescence in the dark/light-adapted state(F_V,_ F_V_’), maximum fluorescence in the dark/light -adapted state(F_M,_ F_M_’), minimum fluorescence in the dark-adapted state (Fo), linear electron transport (LEF), quantum yields of non-photochemical quenching (NPQ), photochemical quenching (qL), yield of photochemistry (Y_II_), yield for dissipation by downregulation (Y_NPQ_), yield of other non-photochemical losses (Y_NO_), extent of change in electrochromic shift on rapid light-dark transition (ECSt), the activity of ATP synthase (gH^+^), and the steady-state rate of proton flux (vH^+^) were measured (Supplemental Methods S13).

### Statistical analysis

One-way ANOVA of fluorescence intensity and phenotypic value was performed using the R software, and significant differences between different groups were determined through Bonferroni correction. Differences were considered statistically significant when *P* < 0.01. The correlation between *PtoNGAL1* and *PtoCUC2* gene expression was determined by correlation coefficient at a significance level of *P*<0.05. Data analysis was performed using the R software.

## RESULTS

### Profiling distinct types of poplar leaves traits

Leaf morphology of poplar is varied among different subgroup of *Populus*. In this study, *P. tomentosa* and *P. simonii* leaves with two markedly different leaf traits were analyzed using the PHMF. PHMF-based features integrate completely different plant morphological features into a single descriptor (Figure 1). In total, 112 phenotypic values were derived from *P. tomentosa* natural population (435 unrelated individuals) and *P. simonii* natural population (505 unrelated individuals), respectively (Supplemental Dataset S1). PCA analysis showed that the top five principal components (PC1–5) explained 61.2% and 72.9% of the phenotypic variation in *P. tomentosa* and *P. simonii*, respectively (Supplemental Figure S2). Among top five principal components, 35 and 27 PHMF-based features with over 2% contribution of leaf phenotypic variation were respectively selected from *P. tomentosa* and *P. simonii* for future analysis. In *P. tomentosa*, 420 of 595 (77.1%) PHMF-derived traits were significantly correlated with each other (*P*<0.01). Nine of 35 (25.7 %) estimated CVs were greater than 50% (Supplemental Dataset S2). Leaf 14, leaf 19, leaf 20, leaf 29, leaf 30, leaf 31, leaf 32, leaf 33 and leaf 34 trait showed *H^2^* values greater than 0.4 (Supplemental Dataset S2). In *P. simonii*, 252 of 351 (71.7%) PHMF-derived traits were also significantly correlated with each other (*P*<0.01). Ten of 27 (37.1 %) estimated CVs of PHMF-derived traits were greater than 50% (Supplemental Dataset S2). Leaf 14, leaf 17, leaf 18, leaf 19, leaf21, leaf 22, leaf 23, and leaf 24 trait showed *H^2^* values greater than 0.4 (Supplemental Dataset S2). Traditional Multivariate Leaf Phenotype (TMLP) data, including LL, LW, LA, LP, AR, AP, and PL, were also obtained in both population (Supplemental Dataset S3). The CV values of seven leaf phenotypes ranged from 10% to 33%. It is worth noting that only average∼ 4.7% PHMF-TMLP trait pairs were significantly correlated (Figure 2A; Supplemental Figure S3), suggesting that the PHMF-derived traits represent novel morphological characteristics of poplar leaves.

**Figure 1.**
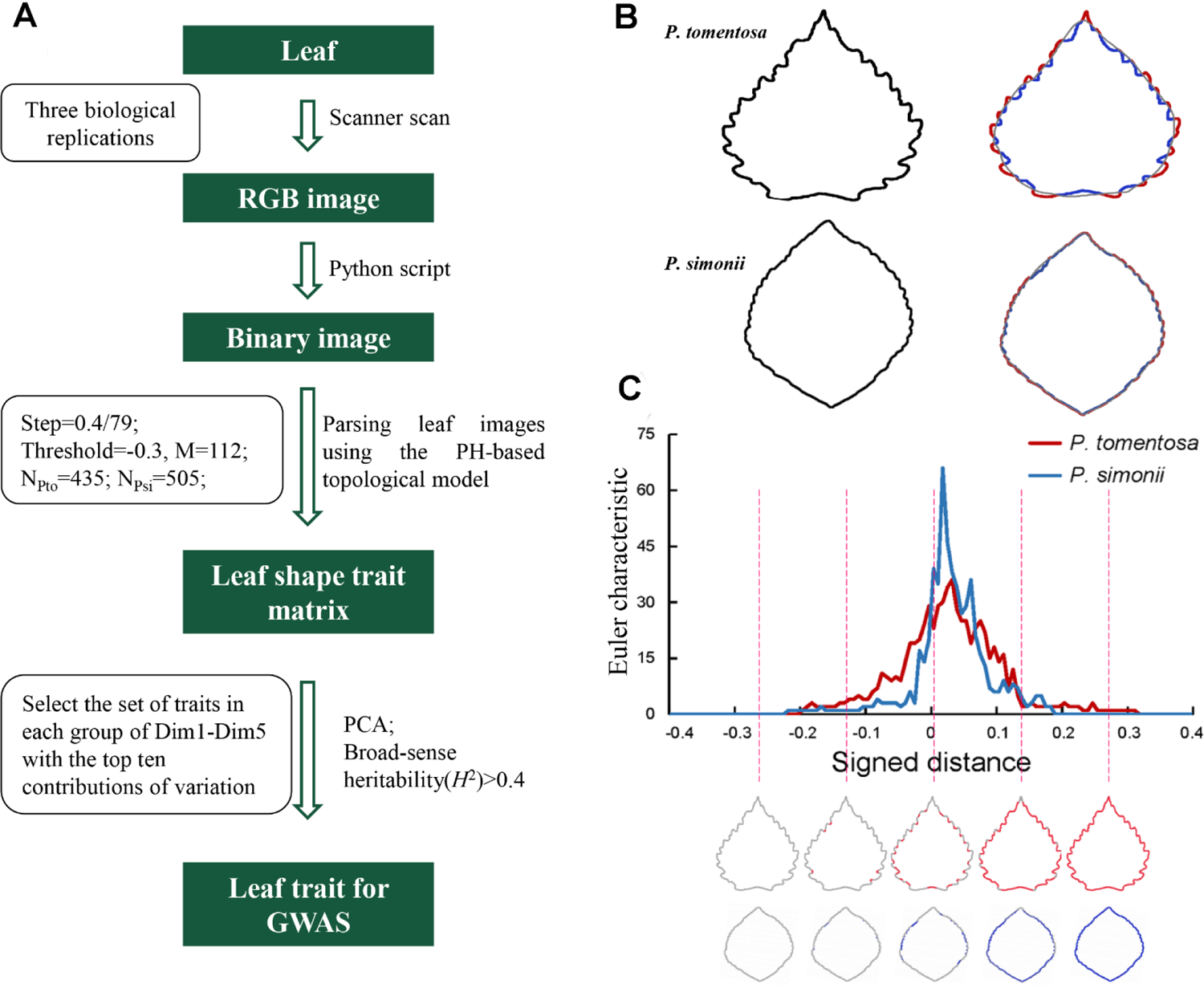
Persistent homology mathematical framework (PHMF) for leaf trait analysis. A, Flowchart showing the genetic basis of leaf morphological characteristics, as revealed by PHMF and topological methods. Important parameters are showed in the box. B, The contour of a *P. tomentosa* and *P. simonii* leaflet. Plot of the signed distance function from the leaf contour to the elliptical Fourier descriptors; values outside the contour are positive (red) and those inside the contour are negative (blue). C, The Euler curve showing the Euler characteristics according to the signed distance function. The number of components (in red) connected at the indicated locations (dashed lines) are shown in the curve.

**Figure 2.**
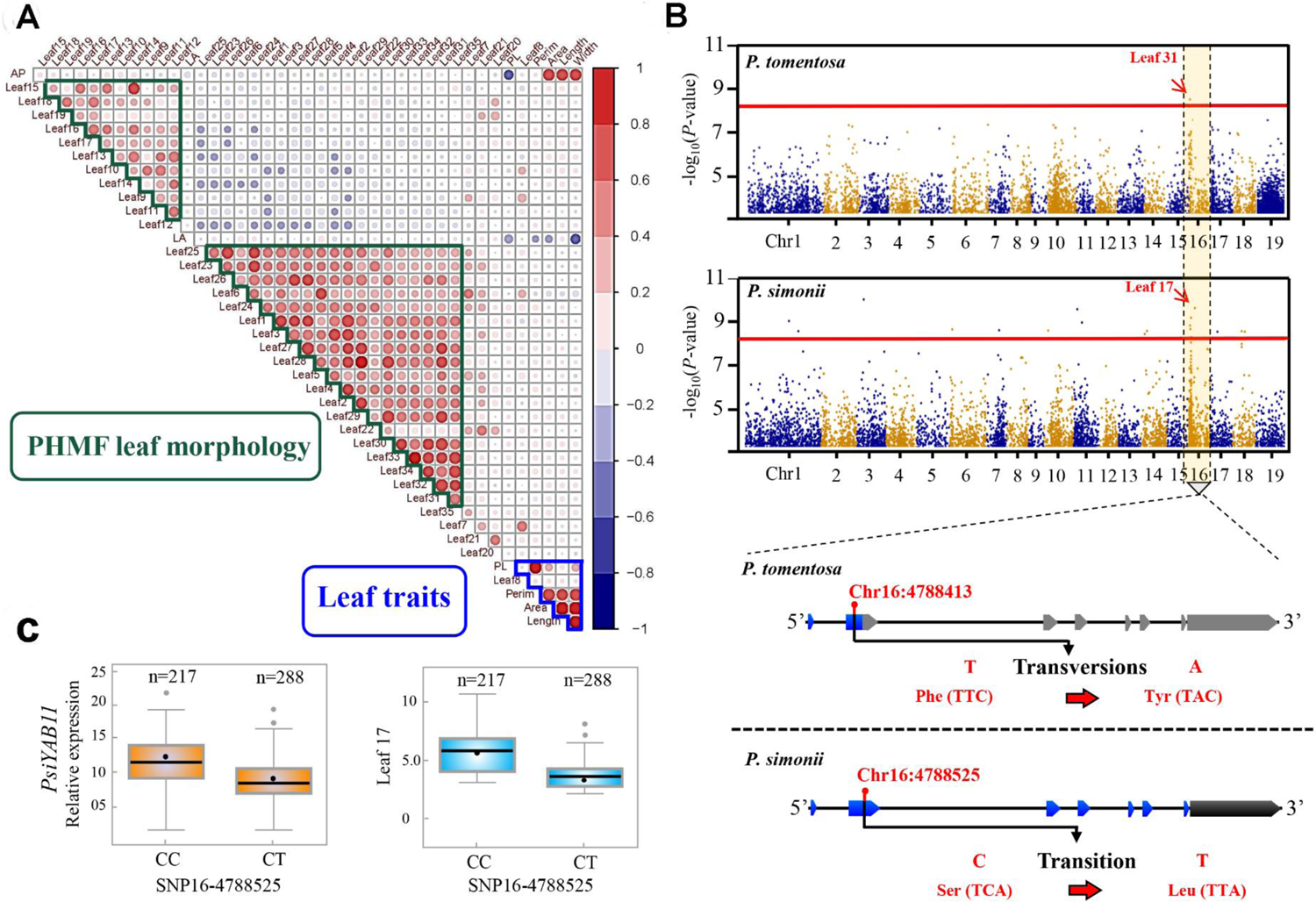
Identification of possible mechanisms underlying natural variation in leaf morphology. A, Correlations of 35 persistent homology mathematical framework (PHMF)-based phenotypic values with 6 “traditional multivariate leaf phenotype” (TMLP) values in *P. tomentosa*. B, Manhattan plot displaying the genome-wide association study (GWAS) results for the PHMF-based phenotypic values in 435 *P. tomentosa* and 505 *P.simonii* unrelated individuals. Nonsynonymous mutation of SNP16-4788413 resulted in the replacement of a phenylanaline (Phe) with an threonine (Tyr) in *P. tomentosa* population. SNP16_4788525 in *PsiYAB11*, the cytosine (C) allele encodes a nonsynonymous mutation, resulting in the replacement of a Sersine (ser) residue with a Leucine (Leu) one in *P. simonii*. Blue represent exon of YAB11. Black represent 3’ untranslated region. Gray represent premature termination of translation. C, Genetic effect of *PtoYAB11* (SNP16-4788413) on leaf traits and transcript abundances. The number of each genotype is given above the graph. The middle line indicates the median. The box shows the 5^th^ to 95^th^ percentiles and the whiskers indicate the interquartile range; dots outside of this are outliers.

### *YAB11* gene SNPs associated with leaf morphology variation in *Populus*

All of PHMF-derived leaf morphology traits with high *H*^2^ (> 0.4) was analyzed using a MLM implemented in TASSEL v5.2. In total, 161 significant associations were detected among 138 SNPs and nine leaf morphology traits (*P* < 1.81E-08) in *P. tomentosa* (Supplemental Dataset S4). The most significant association was to SNP16_4788413 for leaf 31 trait in *P. tomentosa* population (*P* = 1.16E-08) (Figure 2B). SNP16_4788413 located in the second exon of the *YAB11* gene, the Adenosine (A) allele encodes a nonsynonymous mutation, resulting in the replacement of a phenylalanine (Phe) residue with a tyrosine (Tyr). Candidate gene cloning showed that *YAB11* gene carries a premature stop codon in *P. tomentosa* which was defined as *PtoYAB11* with premature stop codon (*PtoYAB11^PSC^*). In *P. simonii* population, eight leaf morphology traits significantly with associated with 171 SNPs (*P* < 7.65E-08). Among the SNP-PHMF leaf morphology pairs, 161 SNPs were associated with a single trait and 10 SNPs showed pleiotropy (Supplemental Dataset S4). The most significant association was to SNP16_4788525 (*P* = 6.39E-10) for leaf 17 trait which also located in the second exon of the *YAB11* gene. SNP16_4788525 in *PsiYAB11*, the Thymidine (T) allele encodes a nonsynonymous mutation, resulting in the replacement of a Sersine (Ser) residue with a Leucine (Leu) one (Figure 2B). SNP16_4788525, the lead SNP significantly associated with leaf 17 trait, showed high transcript abundances of the CC genotype, conferring a higher level of phenotype variation and expression (Figure 2C). Candidate gene cloning showed that *YAB11* gene with complete open reading frame in *P. simonii* which was defined as *PsiYAB11*. To further confirm the reliability of the SNPs identified through resequencing in this region, we randomly selected ten individuals and cloned the *PtoYAB11^PSC^* region from genomic DNA using Sanger sequencing. The high consistency between Sanger sequencing and NGS-based resequencing (Supplemental Figure S4) indicates that the dataset is highly reliable.

### *YAB* gene family organization and phylogeny in *Populus*

GWAS results indicated that *YAB11* gene might be an important regulator of poplar leaf margin variation. *YAB* transcription factors have two conserved domains: a C2C2 zinc finger-like domain and a *YABBY* domain (Supplemental Figure S5). In *P. simonii* genome, the twelve members of the *YAB* gene family were divided into five subgroups: the INNER NO OUTER (INO)-like, CRABS CLAW (CRC)-like, FILAMENTOUS FLOWER (FIL)-like, YAB2-like, and YAB5-like subgroups (Supplemental Figure S6A). The FIL-like subgroup is the largest subgroup, containing *PsiYAB1*, *PsiYAB3*, *PsiYAB4* and *PsiYAB10*. The YAB2-like subgroup contained three members: *PsiYAB2*, *PtsiYAB8*, and *PsiYAB11*. In *P. tomentosa* genome, INO-like, FIL-like, YAB5-like and CRC-like subgroups organization were same conserved in compare with *P. simonii* genome. Only YAB2-like subgroups were divided into two parts that mutated *PtoYAB11^PSC^* fell into a separate clade (Figure 3A). It is suggest that mutation of *PtoYAB11^PSC^* significantly changed YAB2-like subgroup genome organization, implying *PtoYAB11^PSC^* might have undergone functional differentiation in *P. tomentosa*. In both poplar population, three *YAB* gene family members including *YAB4*, *YAB7* and *YAB10* were also associated with five PHFM leaf traits (Supplemental Dataset S4), suggesting other *YAB* gene family members might play synergistic role in leaf morphological variation.

**Figure 3.**
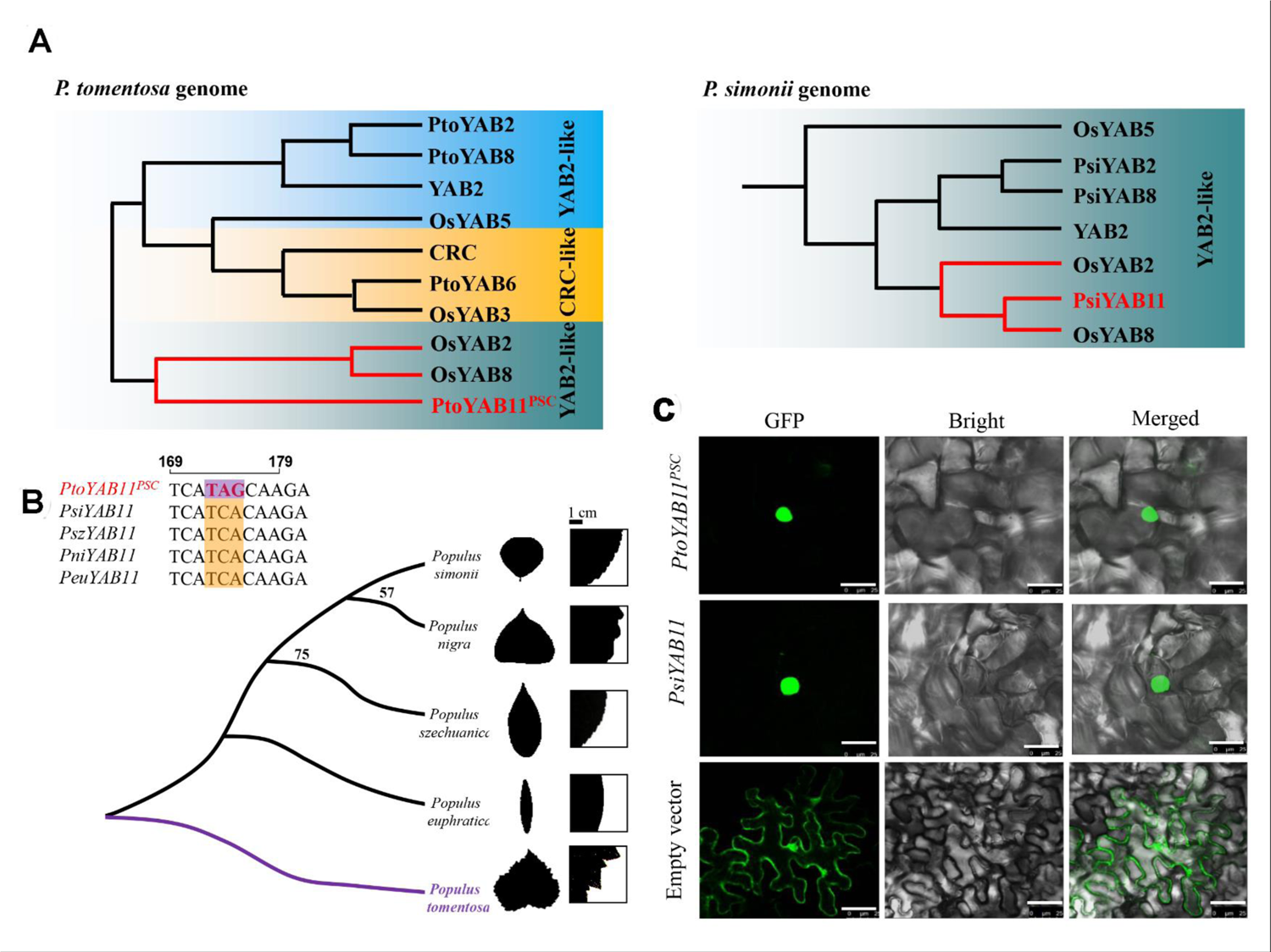
Nucleic acid sequence variation in poplar. A, Phylogenetic analysis of YAB2 group in *P. tomentosa* and *P. simonii* genome. B, Phylogenetic relationships of the poplar *YABBY11* gene. Phylogenetic analysis was performed using the full-length protein sequences of *YABBY11* from four subgenera of *Populus*, including Leuce (*P. tomentosa*), Aigeiros (*P. nigra*), Turanga (*P. euphratica*), and Tacamahaca (*P. simonii* and *P. szechuanica*). Bars, 1 mm. C, Subcellular localization of *PtoYAB11* and *PsiYAB11*. Transient expression of 35S: *PtoYAB11*-GFP and 35S: *PsiYAB11*-GFP in *Nicotiana benthamiana* leaves is shown. Empty vector was used as a control group. Bars, 20 μm.

To determine the evolutionary relationships of *YAB11*, a phylogenetic analysis was performed using the full-length protein sequences of *YAB11* from different subgenera of *Populus*. The phylogenetic dendrogram constructed by the ML method formed two well-defined branches (Figure 3B). One branch contained only *PtoYAB11^PSC^* derived from *P. tomentosa*, and the other branch contained four *YAB11* from *P. simonii* (*PsiYAB11*), *P. nigra* (*PniYAB11*), *P. szechuanica* (*PszYAB11*), and *P. euphratica* (*PeuYAB11*), respectively. Sequence analysis revealed that DNA sequence identity ranged from 72% to 99%. Sequence alignment analysis showed that a stop-gained variant was only identified at the 174 bp region of the second exon of *PtoYAB11^PSC^* (Figure 3B). We further detected the other member of leuce and its hybrid. The results showed that stop-gained variant was also identified in *P. tremula*, *P. alba* and *Populus alba* × *P. glandulosa* (Supplemental Figure S6B), indicating that *YAB11^PSC^* might be leuce-specific variant.

### eQTNs analysis of PtoYAB11^PSC^ in P. tomentosa

To determine whether premature termination of translation affects the transcript regulation function of *PtoYAB11^PSC^*, subcellular localization analysis was used to reveal nuclear localization of *PtoYAB11^PSC^*-GFP and *PsiYAB11*-GFP, respectively. The results showed both PsiYAB11 and truncated protein of PtoYAB11^PSC^ are located in cell nucleus (Figure 3C). Gene expression pattern indicate that both of *PtoYAB11^PSC^* and *PsiYAB11* gene expression were induced by drought, cold, heat and salt stress treatment (Supplemental Figure S7). To further elucidate the regulation network of *PtoYAB11^PSC^* and *PsiYAB11*, *P. tomentosa* and *P. simonii* population gene expression and genetic variants in all genes were subjected to eQTN analyses. In the *P. tomentosa* population, the analyses included 326 common SNPs (MAF > 0.05 and missing rate < 20%) derived from *PtoYAB11^PSC^* and its flanking region (± 2,000 bp). A total of 32,073 gene transcripts were used as the phenotypic variables in the eQTN analysis. A total of 97 unique SNPs were found to be significantly associated with the expression of 55 genes (*P* < 1.5E-04). eQTNs can be divided into *cis*-eQTNs and *trans*-eQTNs according to the distance between the SNP and the gene in question. In total, 42 *cis*-eQTNs were detected for seven genes, and 65 *trans*-eQTNs were detected for 48 genes (*P* < 1.3E-05; Supplemental Dataset S5 and S6). Notably, the expression levels of *PtoYAB4* were negatively associated with *PtoYAB11^PSC^* sequence variation (r=-0.79, *P*<0.01; Supplemental Figure S8), suggesting *PtoYAB4* and *PtoYAB11^PSC^* might exist antagonistic regulatory relationship.

In the *P. simonii* population, the eQTNs analyses included 217 (MAF > 0.05 and missing rate < 20%) common SNPs derived from *PsiYAB11* and its flanking region (± 2,000 bp). A total of 30,193 gene transcripts were used as the phenotypic variables in the eQTNs analysis. A total of 61 unique SNPs were found to be significantly associated with the expression of 52 genes (*P* < 2.3E-04). In total, 27 *cis*-eQTNs were detected for 10 genes and 44 *trans*-eQTNs were detected for 42 genes (*P* < 1.0E-05; Supplemental Dataset S7 and S8). Notably, the most significant associations were between *PsiNGAL1* expression and SNP16_4792605 (*trans*-eQTNs, *P*-value=7.98E-31). For SNP16_4792605, the Thymine (T) allele is the derived allele (A to T), which encodes a nonsynonymous mutation resulting in the replacement of a lysine (Lys) residue with an isoleucine (IIe) one. This mutation changed the sequences of the HMG box element, which is the predicted binding site for *YAB* TFs (Shamimuzzaman et al., 2013). It is suggested that *PsiNGAL1* might be potential downstream targets of *PsiYAB11*.

### Genome-wide binding targets of *PsiYAB11* in *Populus*

Because of premature transcription termination, PtoYAB11^PSC^ lost its C2C2 zinc finger-like domain and a YABBY domain, resulting in the loss of its DNA binding and transcriptional activation capacity. Then, PsiYAB11 were used to further identify potential genome-wide binding targets of YAB11 in poplar using DAP-seq. The average number of total clean sequencing reads were 21,825,083.3. 97.4% reads were mapped on reference genome (Supplemental Table S2). In total, 220 PsiYAB11 binding sites were identified based on optimal threshold and 164 genes were annotated as target genes (Supplemental Dataset S9). The average PsiYAB11 binding sites width was about 287.5 bp. PsiYAB11binding sites were highly centered on target gene transcriptional start sites (TSS) (Supplemental Figure S9) and up to 60.9% overlapped with gene features (Figure 4A). 37.5% PsiYAB11 binding sites were located in promoter region that distance with transcription start site (TSS) 2.0kb. KEGG pathway enrichment analysis of *PsiYAB11* targets showed that “Photosynthesis”, “Oxidative phosphorylation” “Starch and sucrose metabolism” and “Ribosome” pathway were significantly enriched (Supplemental Figure S10). Gene ontology enrichment analysis of *PsiYAB11* targets showed that “electron transport chain” (GO: 0022900, *P*= 6.7e-08) and “transcription regulator activity” (GO:0030528, *P*=2.1E-06) were significantly enriched in molecular function (Supplemental Table S3).

**Figure 4.**
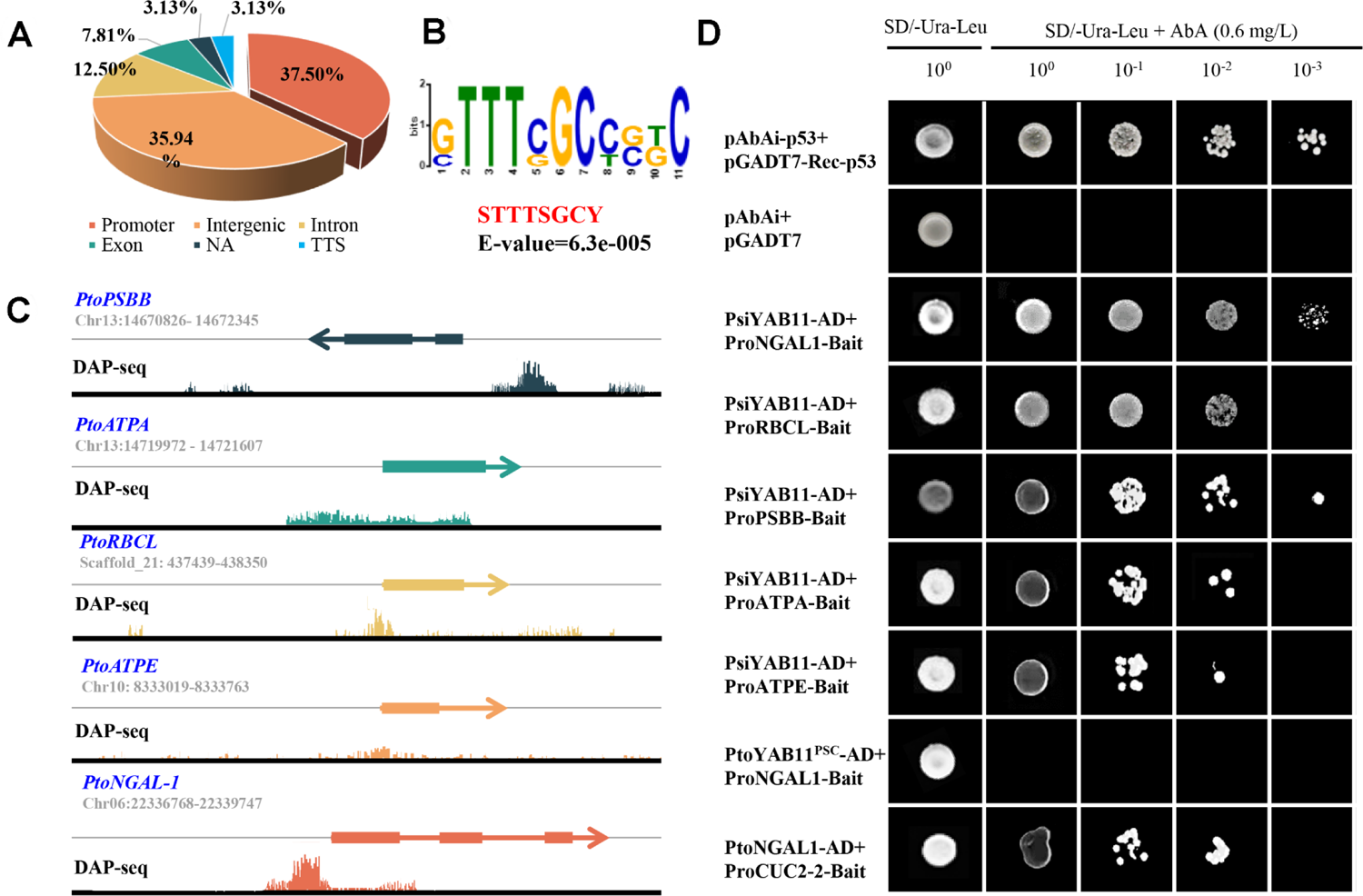
*PtoYAB11 ^PSC^* lost the ability to bind directly to downstream targets. A, The distribution of *PsiYAB11* binding sites relative to gene features. B, Enriched DNA motif identified in *PsiYAB11* binding sites. E-value is shown. C, Integrative genomics viewer (IGV) images of the *PsiYAB11* DAP-seq reads mapping to the promoters of the grapevine *PtoPSBB*, *PtoATPA*, *PtoRBCL*, *PtoATPH* and *PtoNGAL-1* genes. D, Y1H assay showing loss of the ability of *PtoYAB11 ^PSC^* to binding to *PtoNGALs1, PtoPSBB*, *PtoATPA*, *PtoRBCL* and *PtoATPH* promoter fragments. The full-length constructs of *PtoYAB11 ^PSC^ and PsiYAB11* were fused to GAL4 DBD and expressed in the yeast strain AH109 Gold. Transformed yeast was grown in either SD/-Ura-leu-AbA, Trp, or SD/-Ura-leu+AbA media. pGAD-p53+p53-pAbAi and pGADT7-AD+Pro*-PtoNGALs1*–pAbAi were carried out in the same manner as for the positive and negative controls, respectively.

The genomic sequences (<700bp) underlying the binding sites with over 100−log_10_*P* value were used to identify putative motifs bound by PsiYAB11. Three significantly enriched motifs were identified including “STTTSGCY” (E-value=6.3e-005), “AAWSAARAAAAA” (E-value=3.4e-003) and “ACRGGTTAAC” (E-value=1.2e-002) (Figure 4A). To verify the PsiYAB11 binding motif located in their promoter regions, eight potential PsiYAB11 targets were selected to perform DAP-PCR. All binding sites can be amplified from input control and PsiYAB11 DAP samples but not the negative control samples (Supplemental Figure S11), supporting the specificity of the DAP-seq analysis (Figure 4B).

### *PtoYAB11^PSC^* lost transcriptional regulation of *NGAL1*-*CUC2* module in *P. tomentosa*

*NGAL*, *PSBB*, *RBCL*, *ATPA* and *ATPE* genes were identified from eQTNs and DAP-seq analysis results as *PsiYAB11* candidate downstream targets. Y1H assays were used to detect the directly interaction between PsiYAB11 and its downstream targets promoter. Y1H assays provided direct evidence that *PsiYAB11* is specifically bound to the promoters of the *NGAL*, *PSBB*, *RBCL*, *ATPA* and *ATPE* genes *in vivo* (Figure 4B). In contrast, *PtoYAB11^PSC^* lost the regulation of downstream target genes.

Previous studies showed that *NGAL*-*CUC2* module play important role in leaf serrations development. In this study, the expression of *PtoNGAL1* and *PtoCUC2* in the buds, cambium and immature xylem of *P. tomentosa* population was significantly negatively correlated (r = −0.72, *P* < 0.001) (Supplemental Figure S12). Then, *CUC2* were selected as potential targets of NGAL for further Y1H assays. The results also showed that *NGAL* is specifically bound to the promoters of *CUC2* in poplar.

LUC assay was used to validate the transcriptional regulation of *YAB11* and *NGAL1* in poplar. The results showed that both of 35S::*PsiYAB11* and 35S::*PtoYAB4*, co-expressed with *PtoNGAL1* promoters and induced an obvious increase in luminescence intensity (Figure 5A), indicating *PsiYAB11* and *PtoYAB4* are positive regulator of *PtoNGAL1.* In contrast, 35S:: *PtoYAB11^PSC^* co-expressed with *PtoNGAL1* promoters were not significantly changed luminescence intensity, revealing *PtoYAB11^PSC^* had lost its transcriptional activation on downstream targets (Figure 5A). 35S::*PtoNGAL1* co-expressed with *PtoCUC2* promoters significantly decrease in luminescence intensity, suggesting *PtoNGAL1* are negative regulator of *PtoCUC2* (Figure 5B). LUC assay results showed that *PsiYAB11* also positive regulator of *PtoPSBB*, *PtoATPA*, *PtoATPE* and *PtoRCBL* (Figure 5C).

**Figure 5.**
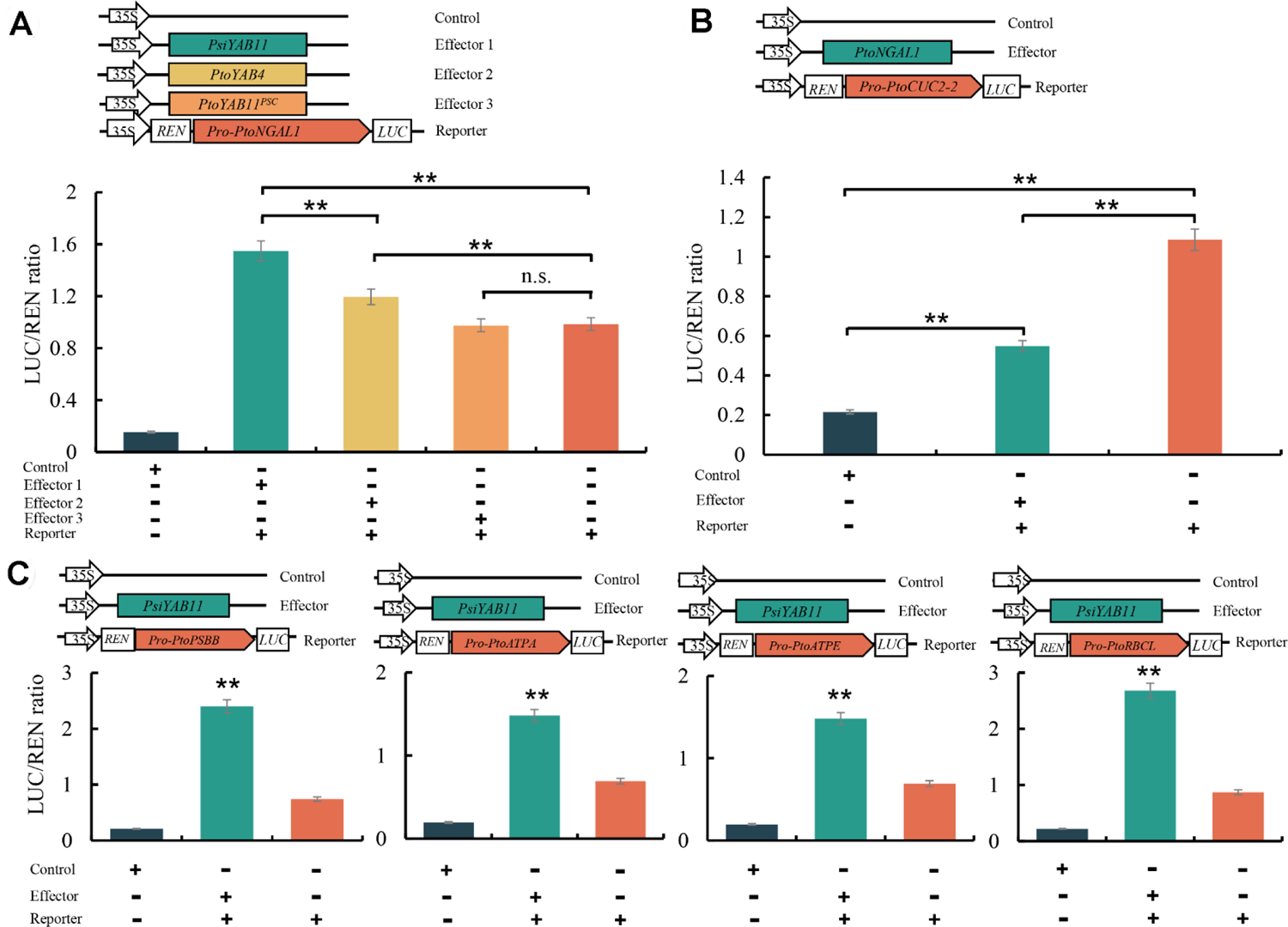
*PtoYAB11 ^PSC^* lost the ability to positive regulate downstream targets. A, Positive regulation of *PtoNGALs1* by *PsiYAB11* was assayed using the dual-LUC system. B, Negative regulation of *PtoCUC2* by *PtoNGAL1* was assayed using the dual-LUC system. C, *PsiYAB11* positive regulate photosynthesis related genes. Data are presented as the means of three biological replicates, and error bars represent SD. Capital letters on error bars indicate significant differences among groups at *P* < 0.01.

### *PtoYAB11^PSC^* promote serrated leaf margins and enhance photosynthesis

To assess the potential biological functions of *YAB11* in the regulation of leaf morphology, transgenic *P. tomentosa* lines overexpressing *PtoYAB11^PSC^* and *PsiYAB11* were generated (Supplemental Figure S1). In total, three *PtoYAB11^PSC^*-overexpressing (OE #11, #19 and #28) and three *PsiYAB11*-OE lines (#4, #11 and #16) were identified (Supplemental Figure S1). Regarding leaf morphology, the LL, LW, LP, and LA of *PsiYAB11-*OE were markedly reduced in comparison to WT. In contrast, *PtoYAB11^PSC^-*OE showed significantly increased LL, LW, LP, and LA and highly serrated leaf margins (Figure 6A, B). *PtoYAB11^PSC^*-OE also showed significantly higher abaxial stomatal size, leaf size, ground diameter and height. Overexpression of *PsiYAB11* in *P. tomentosa* rescued serrated leaf margin development in the WT seedlings, which carry a mutated *YAB11* (Figure 6A). Surprisingly, adaxial stoma also appeared in *PsiYAB11*-OE seedlings (Figure 6C). Overexpression of *PsiYAB11* reduced seeding height, ground diameter, leaf thickness and increased stomatal density (Figure 6D and Supplemental Figure S13). Given the essential role of photosynthesis in leaf and plant development, detailed photosynthesis and fluorescent analyses were performed. *PtoYAB11^PSC^*-OE showed significantly higher Pn, Gs, F_V_/F_M_, Fv/Fo, and F_V_’/F_M_’ values than the other samples. iWUE, LEF, NPQ, and qL were also significantly induced in these *PsiYAB11*-OE samples (Figure 7). Y_II_, Y_NPQ_, Y_NO_, ECSt, vH^+^, and gH^+^ did not significantly differ among the WT, *PtoYAB11^PSC^*, and *PsiYAB11* overexpression samples (Supplemental Table S4).

**Figure 6.**
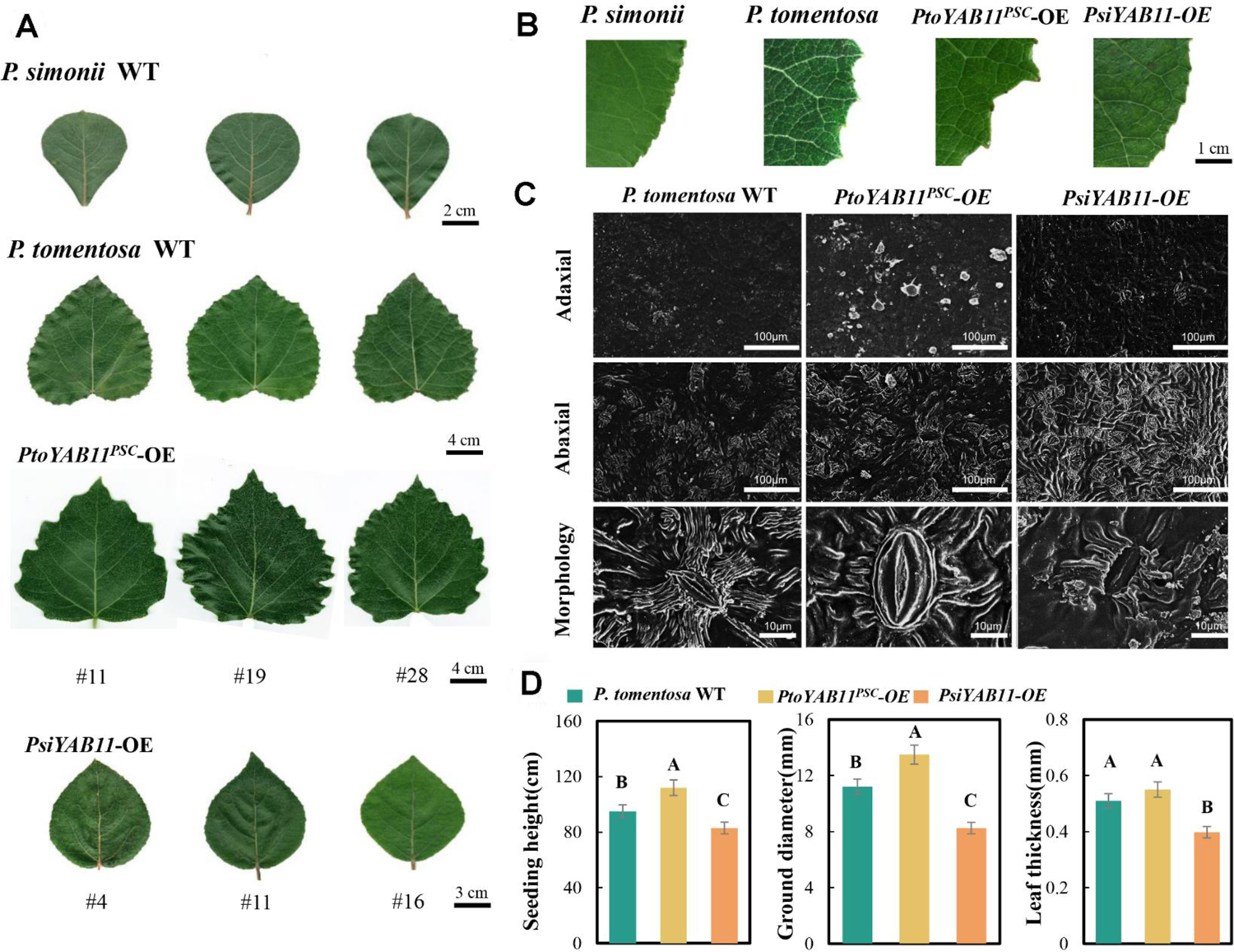
Overexpression of *PtoYAB11 ^PSC^* positively regulates serrated leaf margins and photosynthesis. A, Overexpression of *PtoYAB11 ^PSC^* leading to serrated leaf margins in *P. tomentosa*. B, Detailed view of serrated leaf margins in *PtoYAB11 ^PSC^-*overexpressing (OE) seedlings. C, Stomatal size, and stomatal density in wild-type (WT), *PtoYAB11 ^PSC^-*OE and *PsiYAB11-*OE seedlings. D, Growth traits of wild-type (WT), *PtoYAB11 ^PSC^-*OE and *PsiYAB11-*OE seedlings. Data are presented as the means of three biological replicates, and error bars represent SD. Capital letters on error bars indicate significant differences among groups at *P* < 0.01.

**Figure 7.**
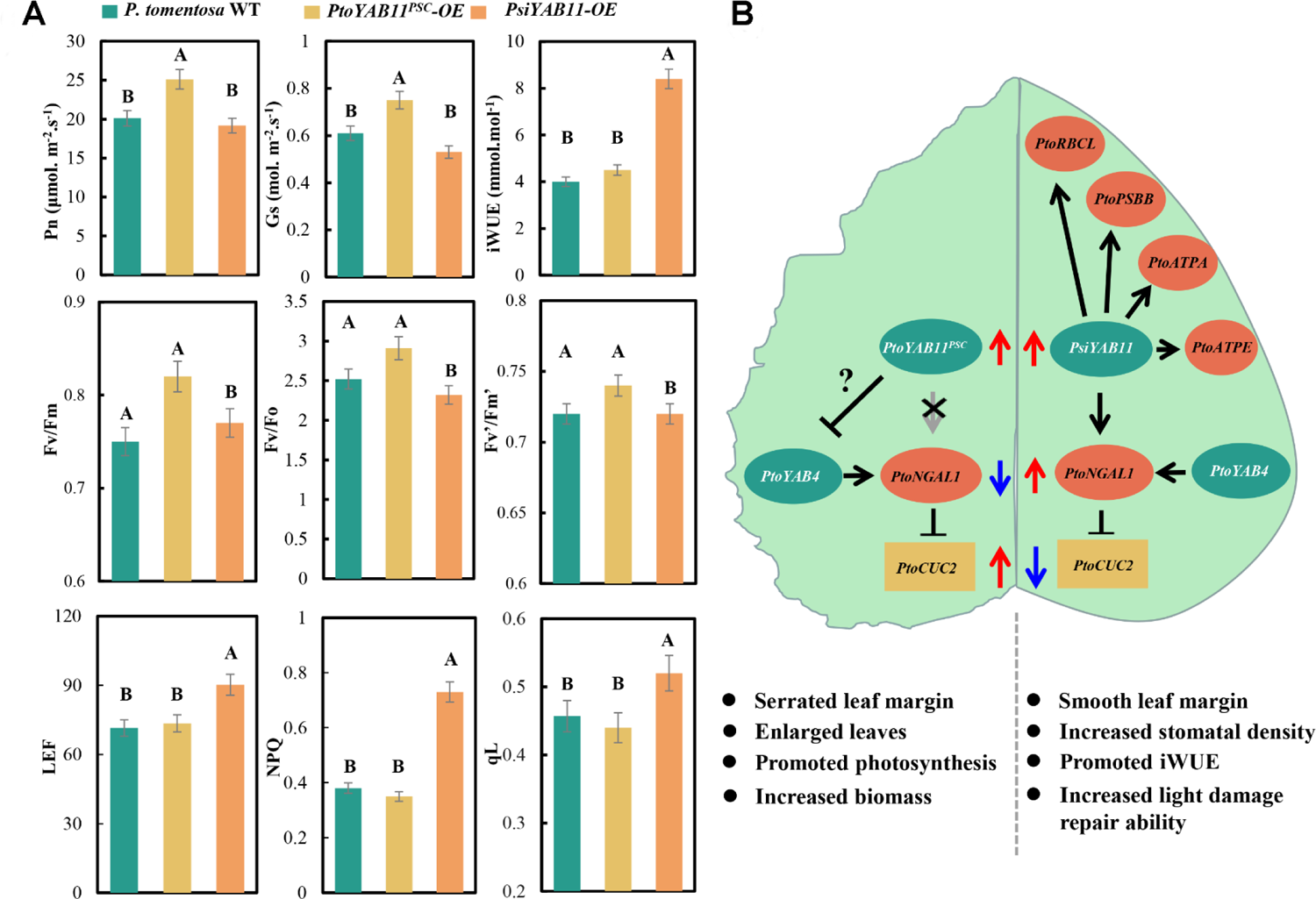
Schematic diagram of *PtoYAB11 ^PSC^*-*PtoNGAL1* module involvement in serrated leaf margins. A, Comparison of photosynthesis and chlorophyll fluorescence parameters among wild-type (WT), *PtoYAB11 ^PSC^-*overexpressing (OE), and *PsiYAB11-*OE *P. tomentosa* seedlings. Pn represents the net photosynthesis rate, Gs represents stomatal conductance, iWUE represents intrinsic water use efficiency, Fv represents variable fluorescence in the dark-adapted state, Fm represents maximum fluorescence in the dark-adapted state, Fo represents minimum fluorescence in the dark-adapted state, Fv’ represents variable fluorescence in the light-adapted state, Fm’ represents maximum fluorescence in the light-adapted state, LEF represents linear electron transport, NPQ represents non-photochemical quenching, and qL represents photochemical quenching (lake model). B, Schematic diagram of the *PtoYAB11 ^PSC^*-*PtoNGAL1* module that leads to serrated leaf margins. Data are presented as mean ± standard error (n = 3). Capital letters indicate significant differences among groups at *P* < 0.01.

## DISCUSSION

### PHMF drive enhanced GWAS of leaf morphology in *Populus*

In previous studies, LW, LA, LL, and LR were measured for genetic regulator mapping analysis (Drost et al., 2015; Chhetri et al., 2019). However, these traits are opposed to multidimensional morphometric analyses of plant morphology and features. Subsequently, Landmarks, pseudo-landmarks, and EFDs have been used to assess plant morphology (Chitwood et al., 2015; Chitwood, 2014; Fu et al., 2016). These high throughput, accurate leaf morphological feature analysis combined with GWAS is greatly improve the analysis accuracy of leaf morphological regulator identification. In this study, 112 phenotypic values representing *Populus* leaf morphology were obtained using PHMF which allowed us to identify previously unknown morphological characteristics of *Populus* leaves. It is fills in the previous cognition of leaf morphology and significantly furthering genotype-phenotype association analysis in *Populus*. However, it provides another major challenge for the future study that how to understand the biological significances of these unknown morphological characteristics. Moreover, although the high throughput, accurate leaf morphological feature analysis is powerful for analyzing leaf morphology, it does not guarantee a positive outcome in core regulator mapping. In *P. tremula*, leaf size and shape metrics obtained by LAMINA were used for further GWAS. The results showed that leaf shape traits of *P. tremula* likely determined by numerous small-effect variations (Mähler et al., 2020). It is worth noting heterochrony and heterophylly are two key features to consider for future study. Heterochrony refers to age-dependent variation in leaf development, while heterophylly may depend on local adaptation. Above all studies indicated that function mapping of multiple locations over many years is of great importance for further candidate gene mapping in relation to leaf morphology.

### *PtoYAB11^PSC^* repressed the *PtoYAB4* transcript abundances promoting serrated leaf margins

*YAB* is a small plant-specific gene family containing a zinc-finger domain in the N-terminal region and a YABBY domain (helix-loop-helix motif) in the C-terminal region (Bowman and Eshed, 2000; Kanaya et al., 2002). *YAB* genes play an important role in lamina outgrowth, maintenance of cell polarity, and establishment of leaf margins (Chen et al., 1999; Watanabe and Okada, 2003; Finet et al., 2016). The FIL-like, YAB2-like, and YAB5-like subgroups, which are expressed in cotyledons, leaves, and floral organs, are considered as significant candidate gene for vegetative development (Golz, 2004; Bartholmes et al., 2012). In this study, *YAB11* was significantly associated with *Populus* leaf margin traits (Figure 2). Premature termination occurred in the second exon of *PtoYAB11^PSC^*, resulting in loss of the zinc finger and *YABBY* domain (Figure 3). Subcellular localization analysis confirmed that *PtoYAB11^PSC^* still located in nuclear (Figure 3D), implying premature termination trimmed the open reading frame (ORF) of *PtoYAB11^PSC^* were still transcribed, translated, and located in the nucleus. Inactive protein-coding genes are typically categorized as pseudogenes (ψs) (Balasubramanian et al., 2009; Xie et al., 2019). Although most of ψs are inactive copies of protein-coding genes, a small fraction of ψs produce regulatory RNAs involved in various fundamental processes (Guo et al., 2009; Wen et al., 2011). Our results showed that *PtoYAB4* activated *PtoNGAL-1* expression and positive regulate leaf 31 traits. Meanwhile, transcript abundance was negatively associated with *PtoYAB11^PSC^* sequence variation, implying *PtoYAB11^PSC^* might be a negative regulator of *PtoYAB4*-*PtoNGAL-1* module. It is suggested that *PtoYAB11^PSC^* not only lost directly regulation on its own downstream targets, but also interferes with the transcriptional regulation network of *PtoYAB4* genes. To determine whether *PtoYAB11^PSC^* could produce regulatory RNA or recruit other biomacromolecules to repress *PtoYAB4* expression, screening of noncoding RNA analysis and yeast double hybrid libraries should be a target of future research.

### *PtoYAB11^PSC^* truncated protein loses its ability to modulate *NGAL1* expression

eQTN mapping is a powerful tool for understanding complex regulatory networks (Zhang et al., 2018). Sequence variations in the gene body and flanking regions inducing polymorphism of transcripts have been defined as *cis*-eQTNs (Hansen et al., 2008). *Trans*-eQTNs are polymorphisms located in the genomic region. A common example of a *trans*-acting eQTL is represented by a TF showing expression polymorphism that regulates downstream transcript targets (Hansen et al., 2008). Our results showed that SNP16_4792605 is associated with the transcript levels of *NGAL1* in *P. simonii* population. SNP16_4792605, located in the HMG box of *PsiYAB11*, has a DNA binding domain that only binds non-B-type DNA conformations (kinked or unwound) with high affinity (Xin et al., 2000). In our study, *PsiYAB11* regulated the expression of its downstream target, *NGAL1*, through its DNA binding domain in *P. simonii*. eQTN mapping, qPCR, Y1H and LUC assays indicated that the *PtoYAB11^PSC^* truncated protein was lost in the HMG box element, leading to a loss of ability to modulate its downstream target *NGAL* in *P. tomentosa*.

In Arabidopsis, *NGATHA1–4* (*NGA1–4*) and *NGAL1–3* TFs belong to the RAV subgroup of the plant-specific B3 family of TFs (Alvarez et al., 2009; Trigueros et al., 2009). *NGALs* act as negative regulators of leaf margin serration. Overexpression of *NGAL3* in Arabidopsis results in smooth leaf margins and “cup-shaped” cotyledons, whereas a loss-of-function allele leads to a serrated leaf margin phenotype (Engelhorn et al., 2012). Recently, more evidences indicate that *CUC2* acts downstream of *NGAL* to promote the formation of leaf margin serrations (Shao et al., 2020). *NGAL1* can directly bind to the *CUC2* promoter, which represses *CUC2* expression (Shao et al., 2020). Repressed *CUC2* or *CUC3* expression leads to smooth leaf margins, suggesting that they promote leaf margin outgrowth and serration formation (Nikovics et al., 2006; Hasson et al., 2011). In the early stages of leaf development, *CUC2* is critical in leaf margin patterns and the initiation of tooth outgrowth (Hasson et al., 2011). In contrast, *CUC3* participates in the maintenance of tooth outgrowth only in later stages (Maugarny-Calès et al., 2019). Our results showed that *PsiYAB11,* with a complete HMG box element, is an upstream regulator of the *NGAL-CUC2* module. In *P. tomentosa*, *PtoYAB11^PSC^* truncated protein lost the ability to promote *NGAL1* expression, leading to increased *CUC2* transcript expression and serrated leaf margins. Furthermore, our results showed that *YAB11^PSC^* are leuce-specific variant. It is suggested that *YAB11^PSC^* might be the primary regulatory factor for leuce-specific leaf margin serration.

### *PtoYAB11^PSC^* bridge environment signaling to leaf morphological plasticity

The *YAB* gene family is reported to not only be involved in diverse developmental processes in plants, but also in responses to various environmental signals. In soybean, 70.5%, 35.2%, and 58.8% of *YAB* members were subjected to drought, NaCl, and ABA treatments (Zhao et al., 2017). Overexpression of *GmYAB10* reduced the survival rate of soybean under PEG and NaCl treatments, indicating that *GmYAB10* is a negative regulator of plant resistance to drought and salt stress (Zhao et al., 2017). Overexpression of *AcYAB4* shortened the root length of Arabidopsis in response to NaCl treatment. This indicates that *AcYAB4* also plays a negative regulatory role in plant resistance to salt stress (Li et al., 2019). Our results showed that both of *PtoYAB^PSC^* and *PsiYAB11* were induced by heat, cold, salt, and drought stress, indicating which were sensitive to environmental signals in poplar. In *Capsella rubella*, *RCO* expression in response to environmental temperature changes results in phenotypic plasticity (Sicard et al., 2014). In this study, *PtoYAB11^PSC^* plays a core regulatory role in *P. tomentosa* leaf margin serration and was also sensitive to various environmental signals, implying that it might be important in the genotype-environment interactions involved in phenotypic plasticity.

Our results showed that *RBCL* genes is downstream targets of *YAB11* in poplar. *RBCL* encode a large subunit of ribulose-1, 5-bisphosphate carboxylase/oxygenase (Rubisco) which play a key role in photosynthetic CO_2_ fixation. Expression of Rubisco large and small subunits should be tightly coordinated to ensure successful assembly. Excessive accumulation of RBCL results in a rapid degradation of the unassembled RBCS, which inhibit chloroplast development (Cohen et al., 2005). In *P. tomentosa*, mutated *PtoYAB11^PSC^* lost positive regulate for *RBCL* gene might benefits for large and small subunits expression tightly coordinated. Our results also showed that YAB11 have function in activated *PSBB*, *ATPA* and *ATPE* gene expression in poplar. These genes involved in light reaction of photosynthesis and played important role in photosynthetic electron and proton transport (Meierhoff et al., 2003). It is explaining that electron transport was only significantly enhanced in ectopic expression of *PsiYAB11* in *P. tomentosa*. In this study, overexpression of *PtoYAB11^PSC^* led to enlarged stomatal size and Gs, which might be the main reason for the iWUE reduction. Moreover, iWUE was determined in various leaf locations that the tip of the leaf has the highest water use efficiency and the base has the lowest (Kirkham et al., 2005). Overexpression of *PtoYAB11^PSC^* affected leaf morphology (LL, LW, perimeter, and leaf margin serration) in our study. We speculated that these changes in leaf morphology also contribute to iWUE regulation, providing new evidence for the adaptive significance of leaf margin morphology.

Based on our findings, we propose a model of action for *PtoYAB11^PSC^* regulate leaf morphology and photosynthesis. In this study, we have provided evidences that *PtoYAB11^PSC^* lost positive regulation of downstream targets *PtoNGAL1* and repressed *PtoYAB4* expression, promoting leaf margin serration. *PtoYAB11^PSC^* also lost positive regulation of light reaction and calvin cycle related genes including *PtoRBCL*, *PtoPSBB*, *PtoATPA* and *PtoATPE*. It is result in *P. tomentosa* photosynthesis induction and light damage repair ability reduction. Hence, we have revealed a new mechanism involved in controlling leaf morphology and provided a candidate module for the genetic improvement of poplar yield and environmental adaptation.

### Accession numbers

All transcriptome expression data (three biological replicates for each group) reported here are available from NCBI with the SRA database accession numbers PRJNA521819, PRJNA521855, PRJNA522886, PRJNA522891, PRJNA357670, SRP141094, SRP073689, and SRP060593. The raw data of genome sequencing have been deposited in the Genome Sequence Archive in the BIG Data Center (BIG, CAS, China) under accession numbers CRA000903 (https://bigd.big.ac.cn/). Euler characteristic curves for leaf serration used in this manuscript can be found at the following GitHub repository: https://github.com/maoli0923/Persistent-Homology-Tomato-Leaf-Root.

## Supplemental data

**Supplemental Figure S1.** PCR validation and gene expression quantitative of wild type, *PtoYAB11*-OE and *PsiYAB11*-OE samples.

**Supplemental Figure S2.** Principal component analysis (PCA) analysis of persistent homology mathematical framework (PHMF)-based leaf morphology features.

**Supplemental Figure S3.** Correlations of 27 persistent homology mathematical framework (PHMF)-based phenotypic values with 6 “traditional multivariate leaf phenotype” (TMLP) values in *P. simonii*

**Supplemental Figure S4.** SNP validation by sanger sequencing.

**Supplemental Figure S5.** The gene structures of YABBY gene family in poplar

**Supplemental Figure S6.** ML tree analysis and sequence alignment of YABBY11.

**Supplemental Figure S7.** Gene expression profiles of YABBY genes in response to different abiotic stress.

**Supplemental Figure S8.** Genetic effect of *PtoYAB11* on *PtoYAB11* and *PtoYAB4* transcript abundances.

**Supplemental Figure S9.** Genome-wide distribution of PsiYAB11 binding sites is centered on gene transcriptional start sites (TSS).

**Supplemental Figure S10.** KEGG enrichment of *PsiYAB11* downstream targets

**Supplemental Figure S11.** DAP-PCR verification of representative binding sites.

**Supplemental Figure S12.** Correlation analysis of PtoNGAL1 and PtoCUC2 gene expression in *P. tomentosa* population.

**Supplemental Figure S13.** Leaf morphology, stomatal size, and stomatal density in wild-type (WT), *PtoYAB11-*OE and *PsiYAB11-*OE seedlings.

**Supplemental Table S1.** Primers used for this study

**Supplemental Table S2.** DAP-seq data mapping information

**Supplemental Table S3.** Gene ontology term enrichment analysis of *PsiYAB11* downstream targets

**Supplemental Table S4.** Phenotype of overexpressed *PtoYAB11* in *P. tomentosa*

**Supplemental Dataset S1.** PH-based leaf shape traits in *Poulus* population.

**Supplemental Dataset S2.** Variable coefficient and broad-sense heritability of PH-based leaf trait

**Supplemental Dataset S3.** Traditional leaf shape traits in *Poulus* population.

**Supplemental Dataset S4.** Association results identified using nine PH-based leaf trait

**Supplemental Dataset S5.** The reported cis-eQTL of *Poulus tomentosa* population

**Supplemental Dataset S6.** The reported *trans*-eQTL of *Poulus tomentosa* population

**Supplemental Dataset S7.** The reported *cis*-eQTL of *Poulus simonii* population

**Supplemental Dataset S8.** The reported *trans*-eQTL of *Poulus simonii* population

**Supplemental Dataset S9.** Annotation of *PsiYAB11* targets

## Acknowledgments

Y.S. conceived and designed the experiments. P.L. carried out the gene expression analysis and drafted the manuscript. D.Z. and M.D. assisted with the writing. P.F., L.Z., and C.B. performed the RNA extractions and participated in the statistical analyses. All authors read and approved the final manuscript.

The English in this document has been checked by at least two professional editors, both native speakers of English. For a certificate, please see: http://www.textcheck.com/certificate/nbawzA

